# Three functionally distinct classes of cGAS proteins in nature revealed by self-DNA-induced interferon responses

**DOI:** 10.1101/2022.03.09.483681

**Authors:** Kenta Mosallanejad, Wen Zhou, Apurva A. Govande, Dustin C. Hancks, Philip J. Kranzusch, Jonathan C. Kagan

## Abstract

Innate immune pattern recognition receptors (PRRs) emerged early in evolution. It is generally assumed that structurally homologous proteins in distinct species will operate via similar mechanisms. We tested this prediction through the study of interferon responses to self-DNA by the enzymatic PRR cyclic GMP-AMP synthase (cGAS). Contrary to expectations, we identified three functional classes of this PRR in mammals. Class 1 proteins (including human) contained a catalytic domain that was intrinsically self-DNA reactive and stimulated interferon responses in diverse cell types. This reactivity was prevented by an upstream N-terminal domain. Class 2 and 3 proteins were either not self-DNA reactive (including chimpanzee) or included proteins whose N-terminal domain promoted self-DNA reactivity (mouse). While self-DNA reactivity of Class 1 cGAS was linked to an ability to access intra-mitochondrial DNA, mitochondrial localization was not associated with other classes. These studies reveal unexpected diversity in the mechanisms of self-DNA reactivity of a PRR.

**One Sentence Summary:** The regulation of self-DNA reactivity of cGAS is evolutionarily diverse in mammals.

## INTRODUCTION

Mammalian cells are equipped with pattern recognition receptors (PRRs) that protect the host through their ability to detect molecular evidence of infection or tissue injury. Among these PRRs is the enzyme cyclic GMP-AMP synthase (cGAS), which synthesizes the second messenger 2’3’-cyclic GMP-AMP (cGAMP) upon binding to double-stranded DNA (dsDNA) in the cytosol (*1*, *2*). As healthy and non-infected cells should not contain cytosolic DNA, the detection of DNA by cGAS is a high-fidelity indicator of infection or cellular dysfunction. cGAS stimulates host defensive responses via the actions of the downstream cGAMP receptor STING (also known as MITA, MPYS, and ELIS). Upon binding cGAMP, STING traffics from the endoplasmic reticulum (ER) to the Golgi apparatus and oligomerizes into a scaffold that serves to activate the kinase tank-binding kinase 1 (TBK1) and the transcription factor IFN regulatory factor 3 (IRF3) (*3*). IRF3 then coordinates the expression of numerous type I interferons (IFNs) and IFN-stimulated genes (ISGs) that promote inflammation and host defense. Although cGAS was initially discovered in the cytosol and therefore regarded as a cytosolic DNA sensor (*1*), recent studies have discovered cGAS in additional subcellular compartments such as the nucleus and the plasma membrane (*4*–*8*). While nuclear cGAS is tightly regulated by proteins in this organelle to prevent chromosomal DNA-induced inflammation (*9*–*14*), DNA in the cytosol binds cGAS and induces conformational changes that drive cGAMP-mediated IFN responses (*15*). Diverse sources of DNA can activate cGAS, including those from infectious agents and host-derived nuclear and mitochondrial DNA (mtDNA) that have reached the cytosol (*16*–*18*). With the exception of subtle difference in the length of DNA detected (*19*–*21*), it is thought that the studies of human cGAS reflect the function of this protein in other mammalian species. However, the symmetry of cGAS functions in nature have largely been explored *in vitro*, where cGAS access to DNA is not an experimental variable. Based on the common *in vitro* behaviors of cGAS, it is somewhat surprising that disparate findings have been made regarding the activities of cGAS within cells—even when studying human cGAS exclusively (*7*, *8*, *22*, *23*).

Human cGAS is a 522 amino acid protein consisting of a basic, unstructured N-terminal domain (1–159 a.a.), and a C-terminal domain (160–522 a.a.) that possesses DNA binding and nucleotidyltransferase (NTase) activities (*1*). Although the N-terminal domain was initially regarded as dispensable for DNA binding and IFN induction (*1*), recent studies have rather reported that the N-terminal domain is required for cGAS functions. For example, several reports indicate that within cells, deletion of the N-terminus of cGAS renders this protein inactive and defective for DNA-induced IFN responses (*7*, *22*, *23*). However, our group found that the deletion of the N-terminus of cGAS leads to IFN responses to self-DNA (*8*). The reason for these disparate datasets is unclear and was explored in detail herein.

In this study, we develop a small molecule activatable genetic system of cGAS-induced self-DNA reactivity. Using this system, we found that the N-terminus of human cGAS prevents the catalytic domain from inducing type I IFN responses against self-DNA in human and mouse cells. We explain previously disparate results on this topic, based on the use of epitope tags that obstruct self-DNA reactivity, and identify amino acids within human cGAS that determine these responses. Evolutionary analysis of self-DNA reactivity revealed three functionally distinct classes of cGAS proteins in nature, with notable differences in self-DNA reactivity being observed in Class 1 (which includes human), Class 2 (mouse), and Class 3 (which includes chimpanzee) cGAS. Leveraging the mechanisms of Class 1 cGAS self-DNA reactivity, we redesigned this PRR to operate as an IFN-inducing sensor of the viral protease activity that is functionally analogous to the naturally occurring guard protein NLRP1 (*24*–*27*). These findings reveal unexpectedly diverse functions of a single PRR in nature, and a means to use synthetic biology to redesign PRR activities in a user-defined manner.

## RESULTS

### Human cGASΔN induces type I IFN responses to self-DNA in human and mouse cells

The functions of the C-terminal DNA binding and catalytic domain of cGAS within cells are debatable. We have reported that within THP-1 monocytes, a human cGAS mutant lacking its N-terminal domain (hcGAS 160–522 a.a., hereafter hcGASΔN) promotes type I IFN expression in the absence of exogenous dsDNA treatment (*8*), whereas others have concluded that hcGASΔN is functionally defective within cells (*7*, *22*). Our prior work relied on the stable expression of hcGASΔN, which drives IFN responses constitutively and may cause aberrant cellular behaviors. To bypass this concern, we devised a distinct genetic system that enables inducible kinetic analysis of cGAS activities. We established a doxycycline (Dox)-inducible expression system for hcGAS full-length (FL) and hcGASΔN in immortalized bone marrow-derived macrophages (iBMDMs). Using this system, we found that Dox-mediated hcGASΔN expression induced the rapid expression of the gene *Ifnb*, which encodes IFN-β (Fig. 1A). In contrast, Dox-induced FL hcGAS expression did not trigger *Ifnb* expression (Fig. 1A). Near-coincident with *Ifnb* expression was the induced transcription of mRNA encoded by the ISG *Rsad2* (radical SAM domain-containing 2) and its product viperin (virus inhibitory protein, endoplasmic reticulum-associated, interferon-inducible) (Fig. 1B and C). Interferon-γ-inducible protein 10 (IP-10), another ISG, was also induced by hcGASΔN (Fig. 1D). The IFN-stimulatory activities of hcGASΔN occurred despite its lower abundance than FL hcGAS within cells (Fig. 1C).

**Fig. 1.**
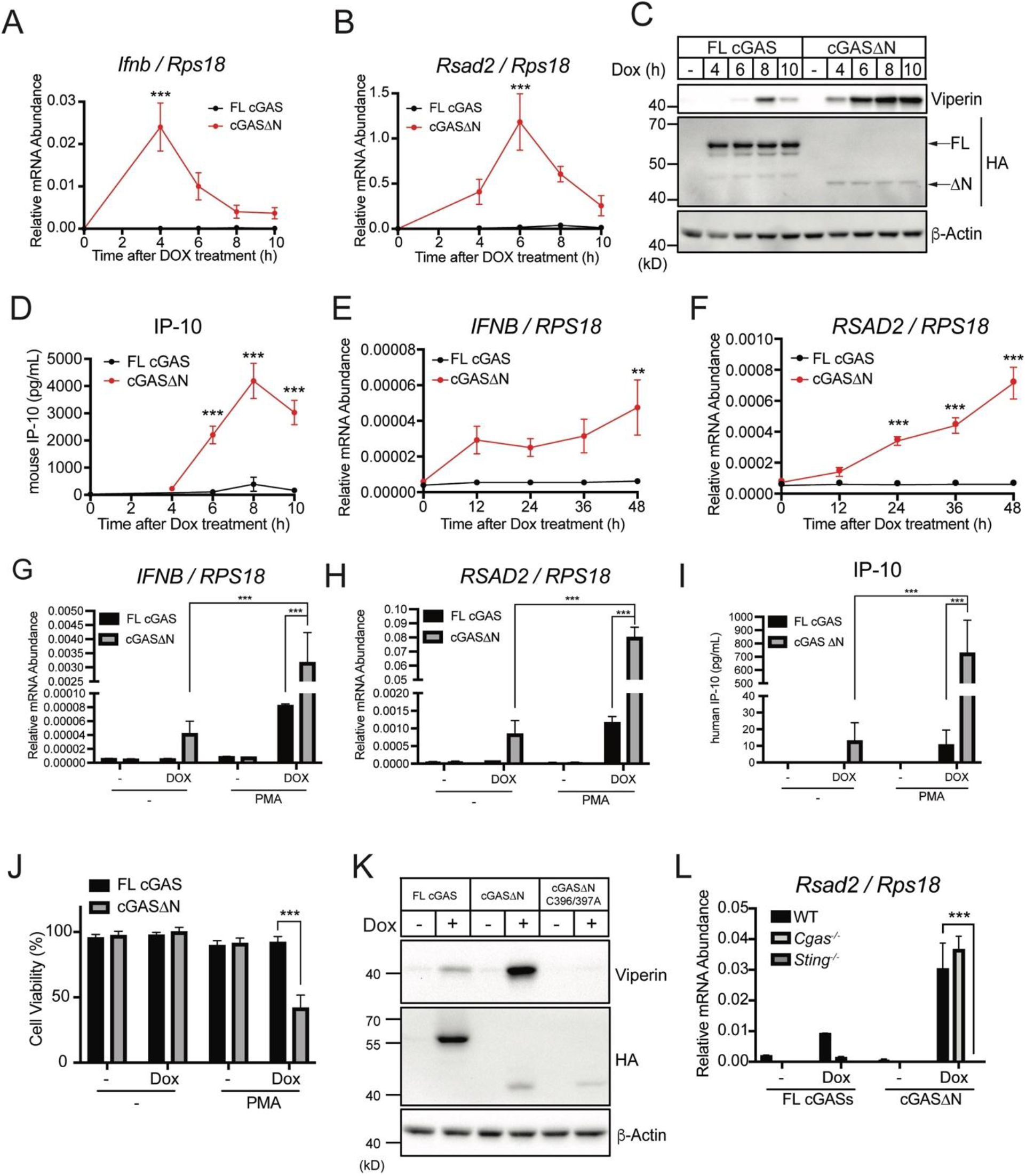
Human cGASΔN induces aberrant type I IFN responses. (**A** and **B**) Real-time quantitative reverse transcription (qRT-) PCR analysis of *Ifnb* (A) and *Rsad2* (B) mRNAs in iBMDMs. Dox was treated to cells for the induction of FL human cGAS or human cGASΔN, and mRNA expression levels were analyzed at the indicated time points. (**C**) Immunoblot analysis of iBMDM lysates after the same treatment as in (A) and (B). Arrows in the HA panel indicate FL human cGAS and human cGASΔN. (**D**) IP-10 ELISA analysis of iBMDM cell culture supernatant of the cells in (A) and (B). (**E** and **F**) Real-time qRT-PCR analysis of *IFNB* (E) and *RSAD2* (F) mRNAs in THP-1 monocytes treated with Dox for indicated time points to induce FL human cGAS or human cGASΔN. (**G** and **H**) Real-Time qRT-PCR analysis of *IFNB* and *RSAD2* mRNAs in THP-1 monocytes treated with Dox together with PMA for 48 hours. (**I**) IP-10 ELISA analysis of THP-1 cell culture supernatant in (G and H). (**J**) Cell viability analysis of THP-1 cells treated as in (G) and (H). (**K**) Immunoblot analysis of lysates from iBMDMs treated with Dox for 8 hours to express FL hcGAS, hcGASΔN, or hcGASΔN with DNA-binding mutation (cGASΔN C396A/397A). (**L**) Real-Time qRT-PCR analysis of *Rsad2* mRNA in WT, *Cgas*^-/-^, or *Sting*^-/-^ iBMDMs treated with Dox for 8 hours to induce expression of FL human cGAS or human cGASΔN. Immunoblot data are the representative from three independent experiments. Graph data are means ± SEM of three (A, B, D, G, H, I, J, and L) or four (E and F) independent experiments. Statistical significance was determined by two-way ANOVA and Tukey’s multicomparison test. Asterisks indicate the statistical significance between FL hcGAS and hcGASΔN at each time point (A, B, D, E, and F) or connected two bars (G, H, I, J, and L). \**P* < 0.05; \*\**P* < 0.01; \*\*\**P* < 0.001.

To validate our results, we applied our Dox-inducible system to THP-1 monocytes. Dox treatment of THP-1 cells containing the hcGASΔN transgene led to the induction of *IFNB* and *RSAD2* mRNAs, whereas cells containing the FL cGAS were poorly immunostimulatory after Dox treatment (Fig. 1E and F). Previously, we found that the treatment of THP-1 cells with phorbol 12-myristate 13-acetate (PMA) potentiated hcGASΔN induced *IFNB* mRNA expression and ultimately caused cell death (*8*). In our Dox system, we observed that PMA enhanced the IFN-inducing ability of hcGASΔN (Fig. 1G-I), with IP-10 production by hcGASΔN increasing over 70-fold after PMA+Dox treatment, as compared to Dox alone (Fig. 1I). This increase in IFN stimulatory activity by PMA correlated with lethality, specifically in cells expressing hcGASΔN (Fig. 1J). The weak immunostimulatory activities of FL cGAS-expressing cells were marginally affected by PMA treatment (Fig. 1G-I). Finally, we applied the same system to normal oral keratinocytes. As a result, keratinocytes treated with Dox induced the expression of *IFNB* and *RSAD2* mRNAs followed by viperin and IP-10 production (fig. S1A-D). Overall, these studies in diverse mouse and human cell types indicate that hcGASΔN triggers type I IFNs that are not induced by FL cGAS.

To determine if hcGASΔN requires DNA binding to induce IFN responses upon expression, we inserted alanine substitutions (C366A/C367A) in hcGAS that abolish DNA binding (*28*). Dox-mediated expression of hcGASΔN C366A/C367A did not induce viperin expression in iBMDMs (Fig. 1K). These data indicate that the C-terminal catalytic domain of cGAS is intrinsically self-DNA reactive, and that this activity is prevented by its upstream N-terminal domain. Further genetic analysis revealed that hcGASΔN did not induce *Rsad2* mRNA or viperin in *Sting^-/-^* iBMDMs (Fig. 1L, fig. S1E), indicating that, as expected, STING is required for hcGASΔN signaling. In contrast, iBMDMs derived from *Cgas*^-/-^ mice induced comparable amounts of *Rsad2* mRNA and viperin protein (Fig. 1L, fig. S1E), indicating that endogenous cGAS is not required for hcGASΔN-mediated signaling. These roles of endogenous cGAS and STING in hcGASΔN-induced IFN expression were confirmed using knockout (KO) cells generated by CRISPR-Cas9 gene editing (fig. S1F-H). Altogether, these data indicated that the N-terminal deletion of human cGAS triggers self-DNA-induced type I IFN responses in a STING-dependent manner.

### Specific amino acids at the N-terminus of hcGASΔN determine self-DNA reactivity

Whereas our data have confirmed that hcGASΔN induces aberrant type I IFN responses, other studies have concluded that N-terminal deletion mutants of cGAS do not elicit IFN expression, even in the presence of exogenous dsDNA (*7*, *22*, *23*). To understand these discrepancies, we noticed that each study has employed different epitope tag positions on their respective hcGASΔN constructs. We have used C-terminally tagged hcGASΔN constructs (*8*), while others have used N-terminally tagged alleles (*7*, *22*, *23*). Therefore, we hypothesized that the different tag positions led to the disparate results regarding hcGASΔN signaling activities. To test this hypothesis, we engineered iBMDMs to express N-terminally or C-terminally hemagglutinin (HA)-tagged hcGASΔN, as well as tag-free hcGASΔN in a Dox-dependent manner. While tag-free hcGASΔN and hcGASΔN-HA induced *Ifnb* mRNA transcripts and ISG expression, HA-hcGASΔN did not induce those responses (Fig. 2A-D). To validate these results, we compared N-terminal or C-terminal green fluorescence protein (GFP)-tagged constructs and found that hcGASΔN-GFP but not GFP-hcGASΔN induced viperin expression (Fig. 2E). Moreover, the addition of an N-terminal FLAG tag to an otherwise active allele (hcGASΔN-HA) inhibited the type I IFN responses induced by this protein (Fig. 2F, G). These results therefore explain reported differences in cGAS function, as the position of the epitope tag determines self-DNA reactivity.

**Fig. 2.**
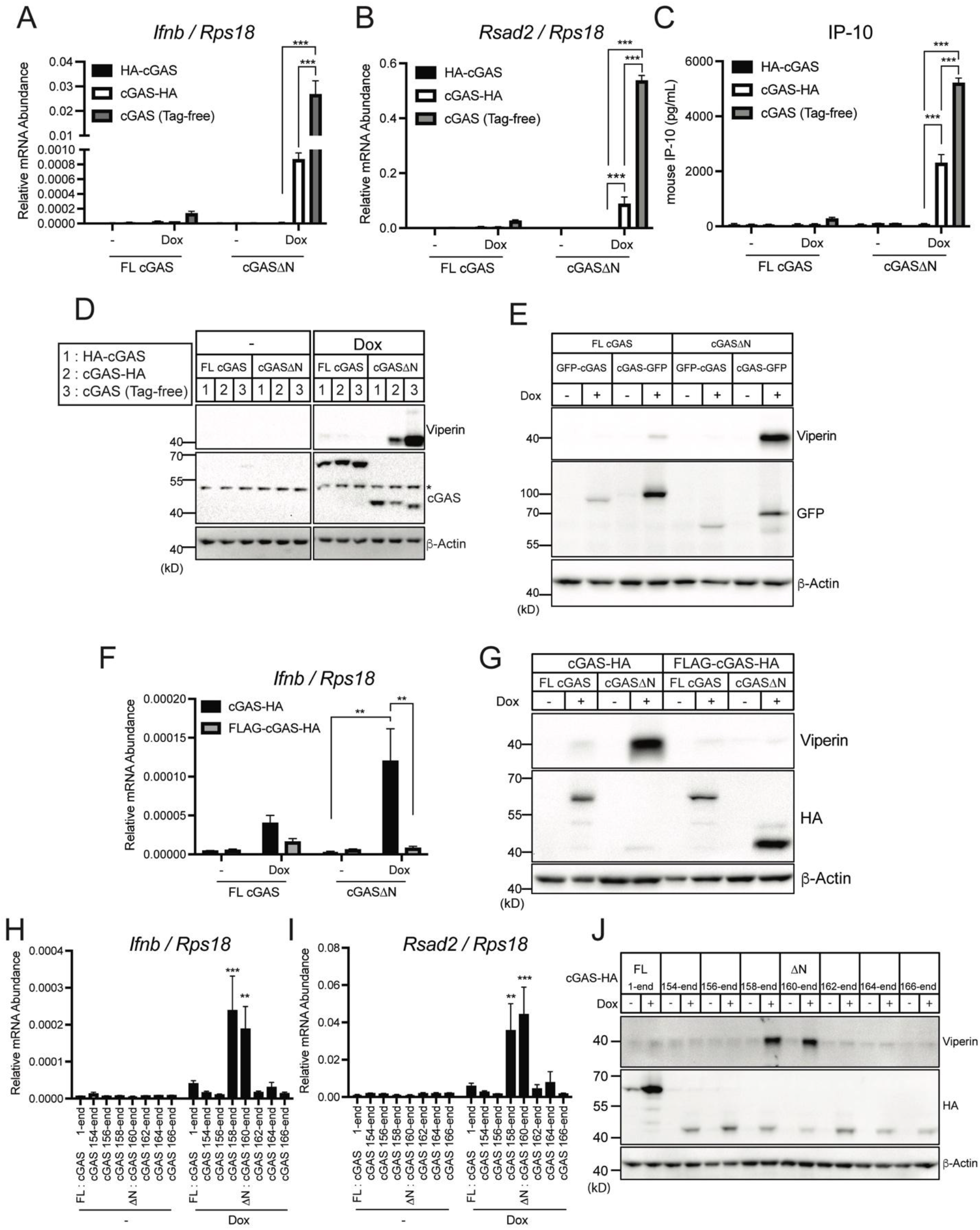
Specific amino acids at the N-terminus of hcGAS ΔN determines self-DNA reactivity. (**A** and **B**) Real-Time qRT-PCR analysis of *Ifnb* (A) and *Rsad2* (B) mRNAs in iBMDMs treated with Dox for 8 hours to induce expression of N-terminally HA-tagged hcGAS (HA-cGAS), C-terminally HA-tagged hcGAS (cGAS-HA), or hcGAS with no tags (Tag-free). (**C**) IP-10 ELISA analysis of iBMDM cell culture supernatant in (A) and (B). (**D**) Immunoblot analysis of iBMDMs treated with Dox as in (A) and (B). The asterisk indicates non-specific bands. (**E**) Immunoblot analysis of iBMDMs treated with Dox for 8 hours to induce expression of N-terminally GFP-tagged hcGAS (GFP-cGAS) or C-terminally GFP-tagged hcGAS (cGAS-GFP). (**F**) Real-Time qRT-PCR analysis of *Ifnb* mRNAs in iBMDMs treated with Dox for 8 hours to induce expression of hcGAS-HA or N-terminally FLAG-tagged hcGAS-HA (FLAG-hcGAS -HA). (**G**) Immunoblot analysis of iBMDMs treated with Dox as in (F). (**H** and **I**) Real-Time qRT-PCR analysis of *Ifnb* (H) and *Rsad2* (I) mRNAs in iBMDMs treated with Dox for 8 hours to induce expression of differently truncated hcGAS. (**J**) Immunoblot analysis of lysates of iBMDMs treated as in (H) and (I). Immunoblot data are the representative from three independent experiments. Graph data are means ± SEM of three (A, B, and F) or five (C, H, and I) independent experiments. Statistical significance was determined by two-way ANOVA and Tukey’s multicomparison test. Asterisks indicate the statistical significance between connected two bars (A, B, C, and F) or between untreated and Dox-treated conditions for indicated hcGAS mutants (H and I). * *P* < 0.05; \*\**P* < 0.01; \*\*\**P* < 0.001.

To further understand the effect of N-terminal tags on hcGASΔN activity, we generated differently truncated hcGAS-HA constructs and expressed each in a Dox-inducible manner in iBMDMs. Among all the hcGAS constructs tested, only hcGASΔN (160–522 a.a.) and hcGAS 158–522 a.a. induced *Ifnb* and *Rsad2* mRNAs, and viperin protein production (Fig. 2H-J). Addition of 4 or more amino acids to the N-terminus of hcGASΔN abolished all signaling activities of this protein. These results indicate that hcGASΔN contains a precise requirement for the N-terminal amino acids for IFN responses to self-DNA.

### Species-specific self-DNA reactivity by mammalian cGAS proteins

The C-terminal catalytic domain of cGAS is highly conserved among mammalian species at the amino acid level, and the functional level *in vitro* (*29*, *30*). We therefore hypothesized that the signaling activity of cGASΔN would be conserved throughout evolution. To assess whether cGASΔN signaling activity is conserved in mice, we expressed mouse cGAS (mcGAS) FL and ΔN in iBMDMs. Whereas some studies have used mcGAS (148–507 a.a.) as the ΔN mutant of mcGAS, other groups have used differently truncated versions (*1*, *23*). Since mcGAS 145–507 a.a. is more similar to hcGASΔN, in terms of the N-terminal primary sequence (Fig. 3A), we tested both versions of mcGAS truncation mutants. Interestingly, both mcGASΔN versions were unresponsive to self-DNA upon Dox-mediated expression in iBMDMs, leading to no IFN activities upon Dox-induction (Fig. 3B-D). FL mcGAS, in contrast, induced strong type I IFN responses upon expression via Dox (Fig. 3B-D). Thus, in contrast to our findings with hcGAS and its ΔN counterpart, mouse FL cGAS is self-DNA-reactive whereas both versions of mouse ΔN are weakly immunostimulatory. These findings prompted a broader evolutionary analysis of self-DNA reactivity by FL cGAS and cGASΔN. We generated iBMDMs that encoded FL and cGASΔN proteins from several non-human primates (NHPs), including orangutan, marmoset, gibbon, chimpanzee, white-handed gibbon, crab-eating macaque, and rhesus macaque (Fig. 3E), and assessed IFN responses upon Dox-induced transgene expression. Like the behaviors of the human proteins, no self-DNA responsiveness was observed for FL cGAS proteins from any NHP examined (Fig. 3F, G). Also similar to human, cGASΔN from orangutan, marmoset, and gibbon all induced *Ifnb* mRNA and viperin protein production, to an extent even greater than what was observed for hcGASΔN (Fig. 3F, G and fig. S2A-D). In contrast, several other NHP cGASΔN proteins, including those from chimpanzee and crab-eating macaque did not trigger type I IFN responses upon expression in iBMDMs (Fig. 3F, G and fig. S2A-D). Most activities observed in iBMDMs were also made in PMA-treated THP-1 cells. For example, hcGASΔN and FL mcGAS induced *IFNB* transcripts, while FL hcGAS and mcGASΔN did not (Fig. 3H). Orangutan cGASΔN was also active in iBMDMs and THP-1 cells (Fig. 3H, I). PMA-mediated potentiation of ISG expression and cell death was also induced in THP-1 cells expressing hcGASΔN, and orangutan cGASΔN, which expressed *IFNB* transcripts (Fig. 3J and fig. S2E-H). However, marmoset and gibbon cGAS were only active in iBMDMs, not in THP-1 cells.

**Fig. 3.**
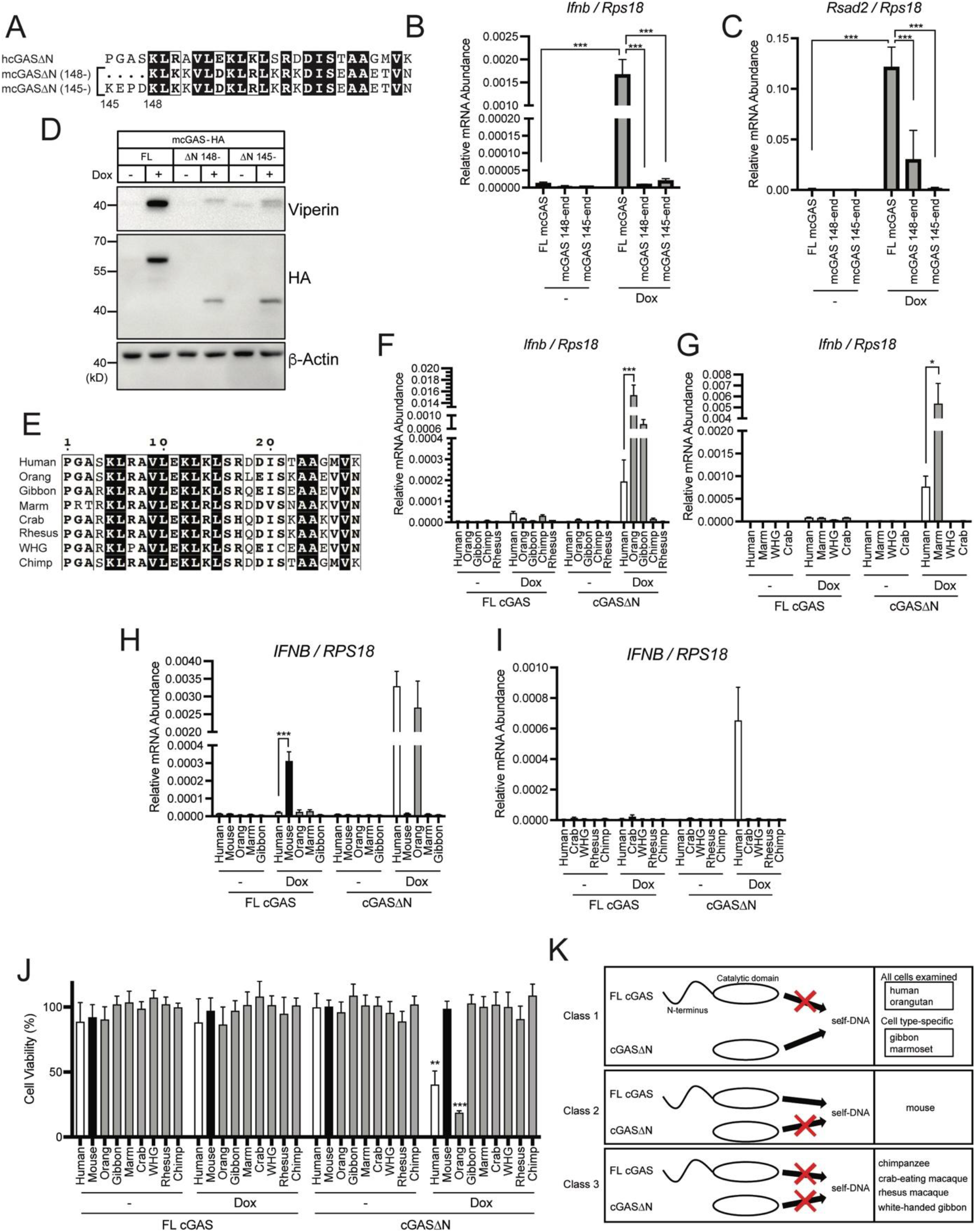
cGASΔN activities are diverse in mammalian species. (**A**) N-terminal amino acid sequences of human cGASΔN, mouse cGAS 145-, and mouse cGAS 148-. (**B** and **C**) Real-Time qRT-PCR analysis of *Ifnb* (B) and *Rsad2* (C) mRNAs in iBMDMs treated with Dox for 8 hours to induce expression of indicated cGAS mutants. (**D**) Immunoblot analysis of iBMDM lysates treated as in (B) and (C). (**E**) N-terminal amino acid sequences of human cGASΔN and non-human primate (NHP) cGAS truncation mutants. Orang: orangutan, Marm: marmoset, Crab: crab-eating macaque, Rhesus: rhesus macaque, WHG: white-handed gibbon, Chimp: chimpanzee. (**F** and **G**) Real-Time qRT-PCR analysis of *Ifnb* (F) and *Rsad2* (G) mRNAs in iBMDMs treated with Dox for 8 hours to induce expression of indicated cGAS mutants. (**H** and **I**) Real-Time qRT-PCR analysis of *IFNB* (H) and *RSAD2* (I) mRNAs in THP-1 monocytes treated with PMA and Dox for 48 hours to induce expression of indicated cGAS mutants. (**J**) Cell Viability of THP-1 cells treated with PMA and Dox as in (H) and (I). (**K**) Model of three classes of cGAS in mammals. In Class 1 cGAS, the N-terminal domain (NTD) inhibits otherwise self-DNA-reactive catalytic domain. In Class 2 cGAS, the NTD promotes and is required for the reactivity to self-DNA. Class 3 cGAS does not react to self-DNA regardless of the presence of the NTD. Immunoblot data are the representative from three independent experiments. Graph data are means ± SEM of three independent experiments. Statistical significance was determined by two-way ANOVA and Tukey’s multicomparison test. Asterisks indicate the statistical significance between connected two bars (B, C, F, G, and H) or between untreated and Dox-treated conditions for indicated cGAS (J). \**P* < 0.05; \*\**P* < 0.01; \*\*\**P* < 0.001

When taken together, we have identified three functional classes of the cGAS proteins in nature. Class 1 proteins restrict self-DNA reactivity by the N-terminal domain in human or mouse cells (or both), and include human, orangutan, gibbon, and marmoset. Class 2 proteins use the N-terminus to promote IFN responses to self-DNA (mouse). Class 3 proteins do not display any self-DNA reactivity (chimpanzee, crab-eating macaque, white-handed gibbon, and rhesus macaque) (Fig. 3K).

### Mitochondrial localization of Class 1 cGASΔN correlates with signaling activity

The different classes of cGAS proteins identified raise questions about the underlying mechanisms of these observations. We considered the possibility that each class of cGAS proteins may display intrinsic differences in DNA-induced cGAMP production. To address this possibility, we incubated recombinant cGAS with DNA *in vitro* and measured cGAMP synthesis. We examined Class 1 human and orangutan cGASΔN, Class 2 mouse FL cGAS, and Class 3 chimpanzee cGAS. The Class 1 and 2 proteins chosen were all active as IFN inducers within cells, whereas Class 3 chimpanzee cGAS was not (Fig. 3F-I). We also included N-terminally FLAG-tagged hcGASΔN and mouse and chimpanzee cGASΔN, all of which were inactive at self-DNA-induced IFN expression within cells. Despite notable differences in self-DNA reactivity associated with each class of proteins within cells, all the cGAS proteins examined behaved similarly in these *in vitro* assays (Fig. 4A). All cGAS proteins were able to synthesize cGAMP in response to synthetic dsDNA, but not to nucleosomal DNA that was isolated from iBMDMs (Fig. 4A). Protease treatment of nucleosomes rendered the resulting DNA capable of stimulating cGAMP production by all classes of cGAS examined (Fig. 4A). The inability of cGAS to produce cGAMP in response to nucleosomal DNA was reported for human cGAS (*6*, *10*–*14*, *31*), but our findings suggest that this inability extends across all three classes of cGAS we have examined. Therefore, the diversity in cGASΔN signaling activities is not due to protein intrinsic activities, but rather by cellular factors that regulate its activation.

**Fig. 4.**
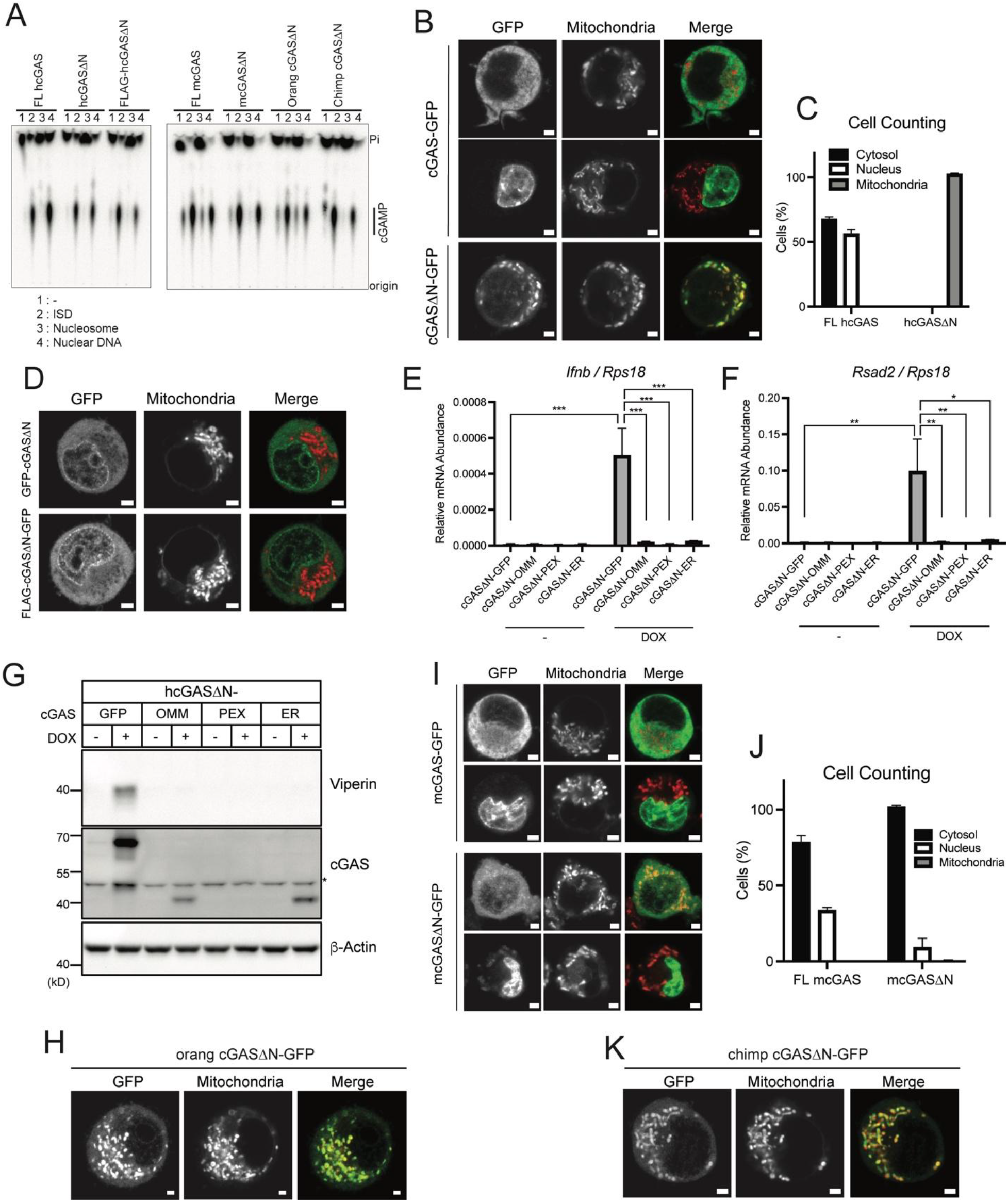
Mitochondrial localization and signaling activities of cGAS. (**A**) cGAS production of cGAMP *in vitro* with purified components. Recombinant cGAS proteins, including FL hcGAS, hcGASΔN (160-522 a.a.), FLAG-hcGASΔN, FL mcGAS, mcGASΔN (148-507 a.a.), orang cGASΔN, and chimp cGASΔN, were activated with indicated double-stranded DNA. cGAMP formation was monitored by incorporation of [ α-^32^P] ATP. Reactions were visualized by treating with alkaline phosphatase and separating by thin-layer chromatography. (**B**) Live confocal micrographs of C-terminally GFP-tagged human FL cGAS or cGASΔN. Representative cell images are shown. (**C**) Cell counting in (B). Cells with nuclear, cytosolic, and mitochondrial cGAS localization were counted and the ratio was calculated. (**D**) Live confocal micrographs of human cGASΔN with indicated tags. Representative cell images are shown. (**E** and **F**) Real-Time qRT-PCR analysis of *Ifnb* (E) and *Rsad2* (F) mRNAs in iBMDMs treated with Dox to induce expression of hcGASΔN with indicated localization tags. (**G**) Immunoblot analysis of iBMDM lysates treated as in (E) and (F). The asterisk indicates non-specific bands. (**H**) Live confocal micrographs of C-terminally GFP-tagged orangutan (orang) FL cGAS or cGASΔN. Representative cell images are shown. (**I**) Live confocal micrographs of C-terminally GFP-tagged mouse FL cGAS or cGASΔN. Representative cell images are shown. (**J**) Cell counting in (I). Cells with nuclear, cytosolic, and mitochondrial cGAS localization were counted and the percentages of each localization pattern were calculated. (**K**) Live confocal micrographs of C-terminally GFP-tagged chimpanzee (chimp) FL cGAS or cGASΔN. Representative cell images are shown. Scale bars in all the images indicate 2 μm. Green signals indicate GFP, and red signals indicate mitochondria in all the merged images. Images and immunoblot data are representative from three independent experiments. Graph data are means ± SEM of three independent experiments. Statistical significance was determined by two-way ANOVA and Tukey’s multicomparison test. Asterisks indicate the statistical significance between connected two bars. \**P* < 0.05; \*\**P* < 0.01; \*\*\**P* < 0.001.

We considered the possibility that access to intracellular DNA may underlie the phenotypes associated with each class of cGAS proteins. To test this idea, we determined the subcellular localization of hcGASΔN by microscopy. Live cell confocal imaging of C-terminally GFP-tagged hcGAS in iBMDMs revealed that hcGASΔN localized in the mitochondria, while FL hcGAS-GFP was distributed in either the cytosol or the nucleus (Fig. 4B and C and Movie S1). Interestingly, when an N-terminal GFP or HA tag was placed onto hcGASΔN, mitochondrial localization was abolished (Fig. 4D). Similarly, N-terminally tagging hcGASΔN abolished IFN activities, as described in Figure 2E. These findings are consistent with a recent study by Chen and colleagues in human cells (*32*). We reasoned that if mitochondrial localization was important for hcGASΔN signaling, forcing its localization to distinct subcellular locations should prevent IFN activities. We therefore engineered hcGASΔN to contain C-terminal membrane localization sequences that direct this protein to the outer mitochondrial membrane (OMM), the peroxisomes, or the ER. This was accomplished by appending onto hcGASΔN transmembrane domains from MAVS (OMM), Pex13 (peroxisomes) or VAMP2 (ER). When expressed via Dox in iBMDMs, none of the membrane-targeted hcGASΔN proteins induced type I IFN responses (Fig. 4E-G). These data indicate that restricting hcGASΔN from access to mitochondria prevents self-DNA reactivity.

To determine if mitochondrial localization was also linked to self-DNA responses induced by other classes of cGAS in nature, we examined the localization of orangutan cGASΔN (Class 1) and FL mcGAS (Class 2), both of which are self-DNA reactive, and the inactive chimpanzee cGASΔN protein (Class 3). Like its human counterpart, orangutan cGASΔN localized to mitochondria, suggesting that IFN activities and mitochondrial localization are a common feature of class 1 cGAS proteins (Fig. 4H). Interestingly, we found that mouse FL cGAS, which is self-DNA reactive, was not localized to mitochondria (Fig. 4I and J) and chimpanzee cGASΔN, which is not self-DNA reactive, was localized to mitochondria (Fig. 4K). Therefore, the correlation of mitochondrial localization and signaling activity of cGASΔN perfectly explains the activities of Class 1 cGAS proteins, but not those of Class 2 or 3. These results suggest that the regulation of self-DNA reactivity for each class of cGAS proteins may differ from each other.

### Viral protease-mediated release of Class 1 cGASΔN induces type I IFN responses

The N-terminal domains of human and mouse cGAS display DNA binding and liquid droplet-forming activities, which are thought to synergize with similar activities in the C-terminal catalytic domain to maximize cGAS responses to DNA (*22*). Studies of Class 2 (mouse) cGAS support this model, as deletion of the N-terminal domain renders cGAS less reactive to DNA *in vitro* (*22*, *23*) and less inflammatory in cells, as compared to FL mcGAS (Fig. 3B-D). However, this theme displays inconsistencies when considering Class 1 human cGAS. cGASΔN from humans is less DNA reactive than FL hcGAS *in vitro* (*22*, *23*), but hcGASΔN is more inflammatory within cells (Fig. 1A-C). The relative contributions of the N- and C-termini to cGAS functions within cells have only been studied in isolation, where cells were engineered to express either of these domains (not both).

We reasoned that if the primary function of the N-terminal domain of cGAS was to promote DNA binding, then the C-terminal catalytic domain would be less functional in the presence of a separate polypeptide encoding its N-terminal domain, as these domains would compete for the same DNA ligands. Alternatively, the mitochondrial localization model would predict that the N-terminal domain could only prevent DNA-induced IFN responses if it was physically attached to the catalytic domain. To test these predictions, we generated a hcGAS transgene that contains a T2A ribosome skip sequence (*33*, *34*) between the N-terminal domain (hcGAS N) and hcGASΔN (Fig. 5A). Thus, a single mRNA would operate as a bicistronic message that produces hcGAS N and hcGASΔN upon translation. We found that Dox-mediated expression of this engineered cGAS within iBMDMs produced hcGASΔN, indicating T2A functionality, and also led to the production of the ISG viperin (Fig. 5B). These results suggest that the N-terminal domain cannot inhibit cGASΔN signaling activity when these domains are separate polypeptides. These data therefore support the idea that the activities present in the C-terminal catalytic domain are sufficient to stimulate IFN responses, and that a central function of the N-terminal domain may be to prevent localization of the C-terminus to self-DNA.

**Fig. 5.**
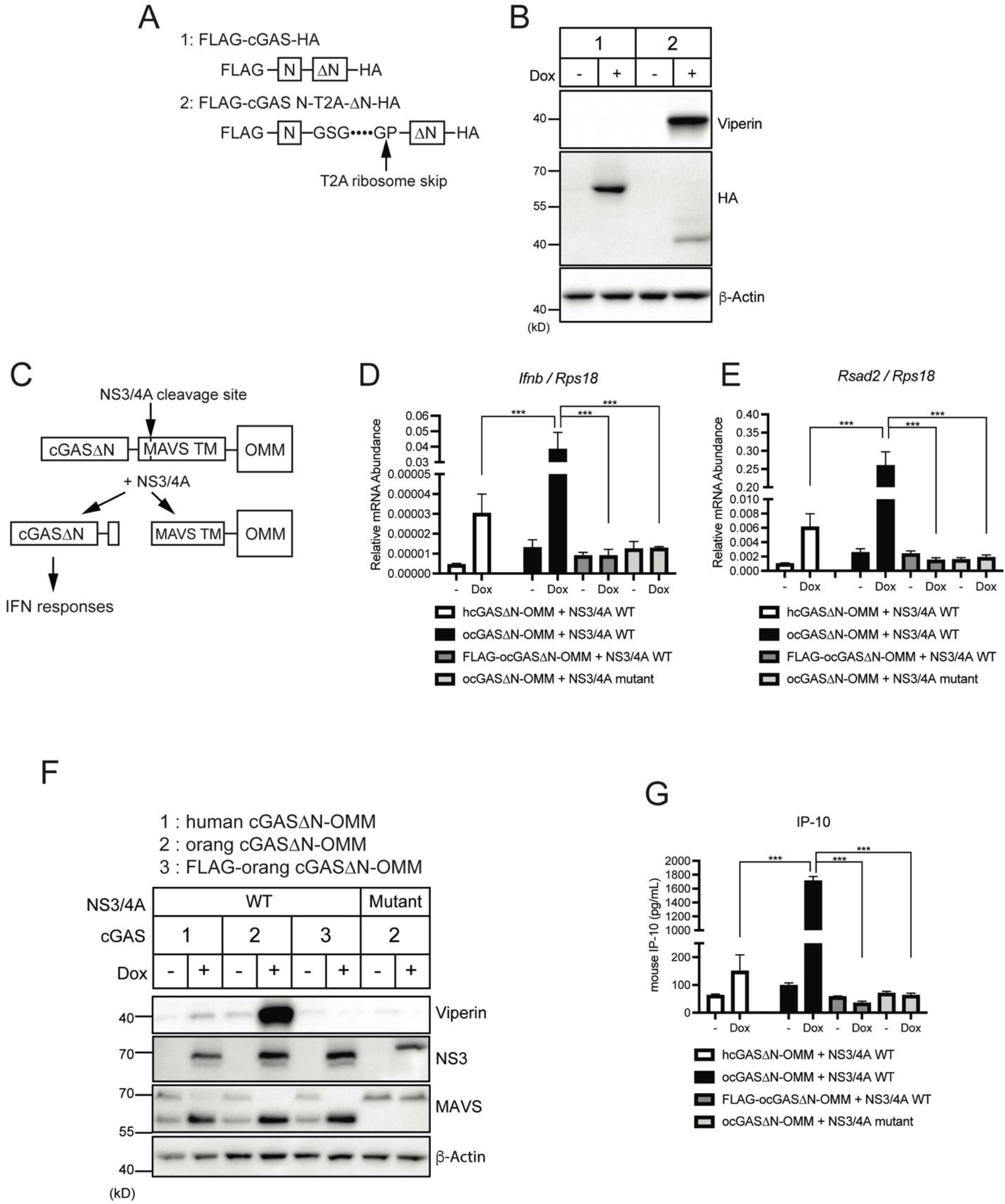
Design of synthetic cGAS as a PRR-guard hybrid that responds to a viral protease. (**A**) Schematic designs of cGAS constructs with T2A ribosome skip consensus sequences. FLAG-cGAS N-T2A-ΔN-HA is separated upon translation at the indicated arrow. (**B**) Immunoblot analysis of lysates of iBMDMs treated with Dox for 8 hours to induce expression of cGAS constructs shown in (A). (**C**) Schematic design of cGAS-outer mitochondrial membrane (OMM) construct and the cleavage by NS3/4A protease. (**D** and **E**) Real-Time qRT-PCR analysis of *Ifnb* (D) and *Rsad2* (E) mRNAs in cGAS-expressing iBMDMs treated with Dox for 24 hours to induce expression of WT or mutant NS3/4A proteases. Cells stably expressed indicated human cGAS (hcGAS) or orangutan cGAS (ocGAS) constructs. (**F**) Immunoblot analysis of iBMDMs treated as in (D) and (E). (**G**) IP-10 ELISA analysis of iBMDM cell culture supernatant in (D) and (E). Immunoblot data are representative from three independent experiments. Graph data are means ± SEM of three independent experiments. Statistical significance was determined by two-way ANOVA and Tukey’s multicomparison test. Asterisks indicate the statistical significance between connected two bars. \**P* < 0.05; \*\**P* < 0.01; \*\*\**P* < 0.001.

Our ability to induce self-DNA reactivity by hcGASΔN, even within cells that contain cGAS N, raised the possibility that other means of dissociating these domains would stimulate IFN production. In this regard, we considered the protein NLRP1, which has emerged as a cytoplasmic sensor of viral and bacterial proteases (*24*–*27*). Cleavage of NLRP1 by pathogen proteases leads to the induction of inflammasome-mediated cell death. In this regard, NLRP1 is considered a guard protein, as opposed to a PRR, with the latter detecting conserved microbial products and the former detecting virulence factor activities (*35*, *36*). No examples of an IFN-inducing sensor of viral protease activity exists. Given that the T2A-mediated separation of the cGAS N- and C-termini was sufficient to trigger IFN responses, it was possible that cGAS can be engineered to operate as an IFN-inducing sensor of viral protease activity. Hepatitis C virus (HCV) is a hepatotropic virus that possesses the protease NS3/4A. This protease is immune-evasive, based on its ability to cleave the RIG-I like receptor (RLR) signaling adaptor MAVS off membranes (*37*). In Figures 4E-G, we anchored hcGASΔN to the outer mitochondrial membrane (OMM) using the transmembrane domain of MAVS, which contains the NS3/4A cleavage site. Therefore, we hypothesized that this hcGASΔN-OMM protein would be cleaved by NS3/4A, leading to the release of hcGASΔN to the cytoplasm and subsequent IFN induction. To test this hypothesis, hcGASΔN-OMM was stably expressed in iBMDMs that encoded a Dox-inducible NS3/4A transgene (Fig. 5C). Expression of hcGASΔN-OMM in the absence of NS3/4A did not lead to any IFN activities, but Dox-mediated induction of NS3/4A stimulated some expression of *Ifnb* and *Rsad2* mRNAs (Fig. 5D, E). Endogenous MAVS in these cells was cleaved upon Dox treatment, confirming the proteolytic function of NS3/4A protein (Fig. 5F). Notably, the induction *of Ifnb and* Rsad2 mRNAs was enhanced substantially when orangutan cGAS was used instead of hcGAS in these assays (Fig. 5D, E). Orangutan cGASΔN-OMM enabled the detection of viperin production and IP-10 secretion upon expression of NS3/4A, which hcGASΔN-OMM barely induced (Fig. 5F, G). Furthermore, and consistent with the importance of mitochondrial localization of Class 1 cGASΔN for self-DNA reactivity, the addition of FLAG-tag to these cGAS constructs abolished IFN responses upon NS3/4A expression (Fig. 5D-G). All Dox-inducible activities were mediated by the protease activity of NS3/4A, as no changes in IFN activities were observed in cells that produced catalytically dead NS3/4A mutant (Fig. 5D-G). These collective data validate the model that the N-terminal domain of Class 1 cGAS proteins prevents self-DNA reactivity and reveal a synthetic biology-based strategy to redesign this PRR into a PRR-guard hybrid, which operates as an IFN-inducing sensor of viral protease activities.

## DISCUSSION

Because the amino acid sequence of cGAS is highly conserved throughout evolution, studies have been largely based on the assumption that the regulatory mechanisms of cGAS in one species hold true for another. In this study, while we confirmed the self-DNA reactivity of the human cGAS catalytic domain, we also identified the evolutionary diversity of cGAS regulation. These results allowed us to stratify the regulation of self-DNA reactivity of mammalian cGAS into three classes (Fig. 3K). Class 1 cGAS in human and the NHPs orangutan, gibbon, and marmoset contains an N-terminal domain that restricts the otherwise self-DNA-reactive catalytic domain. Class 2 cGAS in mouse is the opposite – the N-terminus is required to promote self-DNA reactivity. Class 3 cGAS in other NHPs (chimpanzee, white-handed gibbon, crab-eating macaque, and rhesus macaque) displayed no evidence of an ability to react to self-DNA. This diversity may have derived from unique host-pathogen conflicts in each mammalian species throughout evolution, resulting in differential means of DNA detection and consequently, self-DNA reactivity. Moreover, these observations are intriguing from a clinical perspective, as animal models including mouse, chimpanzee, and rhesus macaque are often components of therapeutic development pipelines.

Our mechanistic analysis revealed an exquisite sensitivity of the N-terminus of human cGASΔN for self-DNA responsiveness, as adding greater than two amino acids onto the N-terminus was sufficient to abolish IFN activities. These findings may be considered from the perspective of the common use of N-terminal epitope tags to study FL and cGASΔN functions, which we now consider to be the cause of much conflicting literature. Future studies should consider the position of the epitope tags to be as important an experimental variable as the species of the cGAS protein under investigation.

Our finding of hcGASΔN localization to mitochondria is consistent with a recent study reporting that human cGASΔN triggers IFN responses upon localization to these organelles (*32*). While these findings explain the operation of human cGAS and other Class 1 cGAS proteins, we found that cGAS proteins in other classes do not follow this rule of self-DNA reactivity. Indeed, FL mcGAS (Class 2) induces IFN responses without localizing to mitochondria and chimpanzee cGASΔN (Class 3) does not trigger these responses even though it localizes to mitochondria. Given that cGAS proteins in all three classes react with DNA and synthesize cGAMP *in vitro*, these findings emphasize the diversity of cGAS regulation that is best revealed from studies within cells. The rules that govern cGAS in Class 2 and Class 3 activities were not revealed in this study and warrant future investigation.

Finally, our redesign of two Class 1 cGAS proteins into PRR-guards that can use self-DNA reactivity as a functional output of viral protease detection is noteworthy. Viral proteases, in particular NS3/4A, are naturally immune-evasive because they cleave host signaling proteins to inactivate PRR-induced responses to infection. Classic therapeutic strategies to target viral proteases involve inhibition, which results in selective pressure for viral escape mutants. Rather than inhibiting viral proteases, our redesigned cGAS has forced the normally immune-evasive NS3/4A protease to operate as an immunostimulant, which offers several opportunities for further study in the context of infection. These findings provide a mandate to consider synthetic biology-based strategies to alter the host-pathogen conflicts that determine infectious outcome.

## MATERIALS AND METHODS

### Study design

The aim of this study was to investigate the self-DNA reactivity of cGAS and its domains in human and other mammals. We investigated the activities of these cGAS and mutants using doxycycline-mediated transient expression system in mouse and human macrophages. Sample sizes for each experiment are indicated in the figure legends.

### Cell culture

Immortalized bone marrow-derived macrophages (iBMDMs), HEK293Ts, and Plat-GP cells were cultured in DMEM (Gibco) supplemented with 10% FBS (Gibco), referred to as complete DMEM, at 37°C in 5% CO_2_. For passage, iBMDMs were lifted using PBS (Gibco) supplemented with 2.5 mM EDTA (Invitrogen) and plated at dilution 1:10. HEK293T and L929 cells were grown under the same conditions as iBMDMs but were passaged by washing with PBS and lifting with 0.25% Trypsin-EDTA (Gibco) with a 1:10 dilution. THP-1 cells were grown in suspension culture using RPMI-1640 media (Lonza) supplemented with 10% FBS, referred to as complete RPMI-1640, at 37°C in 5% CO_2_. For passage, cells were split at a dilution of 1:5. For experiments examining the effects of PMA-induced differentiation of THP-1 cell lines, cells were treated for the 72 hours with PMA (MilliporeSigma) at a concentration of 50 ng/mL. Normal oral keratinocytes were cultured in keratinocyte SFM (Gibco) supplemented with Human Keratinocyte Growth Supplement (Gibco) and passaged by washing with PBS and lifting with 0.25% Trypsin-EDTA with a 1:10 dilution.

### Generating cells with stable or doxycycline-inducible gene expression

cDNAs of wild type and truncation mutant cGAS were amplified by polymerase chain reaction (PCR) using oligonucleotide primers containing restriction enzyme digestion sites. For the amplification of non-human primate (NHP) cGAS, cDNAs in the pcDNA6 vector (*38*) were used as PCR templates. For Dox-induced gene expression, cGAS cDNAs were inserted in the BamHI and NotI restriction sites in pRetroX-TRE3G (TakaraBio) using In-Fusion Snap Assembly Master Mix (TakaraBio). For stable gene expression, cGAS DNAs were inserted in pLenti-CMV-GFP-Puro in replacement of GFP using XbaI and SalI. For fusion mutants and the T2A insertion mutant of cGAS cDNAs, In-Fusion Snap Assembly Master Mix was used to incorporate cGAS and the other fragments together in the vectors. All constructs generated here were sequence-confirmed by Sanger sequencing.

To generate lentiviral particles for the stable expression of transgenes, HEK293T cells were transfected with the packaging plasmids psPAX2 and pCMV-VSV-G along with the transgene in pLenti-CMV-GFP-Puro using Lipofectamine 2000 (Invitrogen). pCMV-VSV-G was a gift from Bob Weinberg (Addgene plasmid # 8454; http://n2t.net/addgene:8454; RRID:Addgene_8454) (*39*). psPAX2 was a gift from Didier Trono (Addgene plasmid # 12260; http://n2t.net/addgene:12260; RRID: Addgene_12260). pLenti CMV GFP Puro (658-5) was a gift from Eric Campeau & Paul Kaufman (Addgene plasmid # 17448; http://n2t.net/addgene:17448; RRID: Addgene 17448) (*40*). All genes of interest were subcloned into the GFP site. Plasmids were transfected into 10 cm^2^ dishes of HEK293Ts at 50–80% confluency using Lipofectamine 2000 by mixing DNA and Lipofectamine 2000 ratio at 1:2. Media was changed on transfected HEK293Ts 16–24 hours after transfection, and virus-containing supernatants were harvested 24 hours following the media change. Viral supernatants were passed through a 0.45 μm filter to remove any cellular debris. Filtered viral supernatants were mixed with 5 μg/mL polybrene (MilliporeSigma) and placed directly onto target cells, followed by the spinfection (centrifugation at 1,250 × g, 30°C for 1 hour). Cell culture media were replaced with the appropriate complete media and cells were incubated for 24 hours. Spinfection with the viral supernatants was repeated on the following day, and cells were used for indicated assays.

Doxycycline (Dox)-inducible gene-expressing cell lines were generated using Retro-X™ Tet-On 3G Inducible Expression System (Takara Bio). Plat-GP cells were transfected with the pRetroX-Tet3G plasmid together with the packaging plasmid pCMV-VSV-G using Lipofectamine 2000. Using viral supernatant of Plat-GP cells, iBMDMs or THP-1 cells were subjected to spinfection in a similar manner as above. After consecutive spinfections for two days, Tet3G-transduced cells were selected using G418 (Invivogen) and single cell clones were isolated. Tet3G-containing cell clones were transduced with TRE3G virus prepared from Plat-GP cells transfected with pRetroX-TRE3G containing genes of interest and packaging plasmid pCMV-VSV-G. TRE3G-transduced cells were selected using puromycin (Gibco) and pooled cell culture was used in each assay. Cells were treated with 1 μg/mL Dox (MilliporeSigma) were subjected to each analysis at the indicated time points in each figure legend.

### Real-Time quantitative reverse transcription (qRT-) PCR

RNA was isolated from cells using Qiashredder (QIAGEN) homogenizers and the PureLink Mini RNA Kit (Life Technologies) and treated with subsequently DNase I (Invitrogen) to remove genomic DNA. Relative mRNA expression was analyzed using the TaqMan RNA-to-Ct 1-Step Kit (Thermo Fisher Scientific) with indicated Taqman probes (Thermo Fisher Scientific) on a CFX384 Real-Time Cycler (Bio-Rad Laboratories). Each C_T_ value was normalized with the mRNA expression of the control genes (*RPS18* for human and *Rps18* for human) and the relative mRNA abundance was calculated by the ΔΔC_T_ method. Taqman probes used in this study are as follows: *Ifnb* (mouse): Mm00439552_s1, *Rsad2* (mouse): Mm00491265_m1, *Rps18* (mouse): Mm02601777_g1, *IFNB* (human): Hs01077958_s1, *RSAD2* (human): Hs00369813_m1, *RPS18* (human): Hs01375212_g1.

### Cell viability assay

Cell viability was measured using CellTiter-Glo (Promega Corporation), a luminescent assay for ATP in living cells. Untreated cells were used as a positive control for 100% cell viability and subjected to serial dilution for the standard curve. Luminescent outputs were read on a Tecan plate reader and viability was calculated using the standard curve.

### Immunoblotting and ELISA analysis

Cells were lysed with RIPA buffer (50 mM Tris-HCl, pH7.5, 150 mM NaCl, 1% TritonX-100, 0.5% Sodium Deoxycholate, 0.1% sodium dodecyl sulfate (SDS), which was supplemented with cOmplete, Mini, EDTA-free Protease Inhibitor Cocktail [Roche] before use) and the lysates were centrifuged at 4°C, 16,000 × g for 10 minutes. Supernatants were mixed with × SDS sample buffer supplemented with Tris(2-carboxyethyl)phosphine hydrochloride (TCEP, Thermo Fisher Scientific) and boiled at 100°C for 5 minutes. Samples were separated by SDS-PAGE and transcribed to PVDF membrane by Immunoblotting. PVDF was blocked with 5% skim milk for 1 hour and probed with indicated primary antibodies over night at 4°C, followed by secondary antibodies (1:2000 dilution) for 1 hour. Primary antibodies used in this study include viperin (1:1000 dilution, MilliporeSigma), HA (1:1000 dilution, MilliporeSigma), b-Actin (1:1000 dilution, Cell Signaling Technology), cGAS (1:1000 dilution, MilliporeSigma), STING (1:1000 dilution, Cell Signaling Technology), and NS3 (1:1000, GeneTex). Secondary antibodies for human, mouse, and rat immunoglobulins (IgGs) were purchased from Rockland Immunochemicals. Culture supernatant from treated cells were collected and applied to the IP-10 antibody-coated plate for ELISA analysis (R&D Systems) following the manufacture’s instruction.

### In vitro 2’3’-cGAMP assay

FL cGAS and cGASΔN recombinant proteins were prepared as previously described (*21, 41*). Briefly, cGAS mutants were cloned into a custom pET vector for expression of an N-terminal 6 x His-SUMO2 fusion protein in *E. coli. E. coli* BL21-RIL DE3 (Agilent) bacteria harboring a pRARE2 tRNA plasmid were transformed with a pET cGAS plasmid, and 6 × His-SUMO2-cGAS recombinant proteins were purified from clarified *E. coli* lysate by binding to Ni-NTA (QIAGEN) and gravity chromatography. The His-SUMO2 tags were removed by dialyzed overnight at 4°C in dialysis buffer (20 mM HEPES-KOH pH 7.5, 300 mM NaCl, 1 mM DTT) after supplementing ~250 μg of human SENP2 protease (fragment D364–L589 with M497A mutation).

Nucleosomal DNA was isolated from untreated iBMDMs using EpiScope Nucleosome Preparation Kit (TakaraBio) and the counterpart naked double-stranded DNA (dsDNA) was obtained by removing histones using Proteinase K following manufacture’s instruction.

cGAS activation and cGAMP synthesis was performed in vitro using purified components and measured with thin-layer chromatography as previously described (*41*). Briefly, 1 μM cGAS recombinant proteins were incubated with the DNA above or 45 bp interferon stimulatory DNA (ISD) (*42*) in the 20 mL reaction buffer containing 50 mM Tris-HCl pH 7.5, 35 mM KCl, 5 mM Mg(OAc)_2_, 1 mM DTT, 25 mM ATP, 25 mM GTP, and [α-^32^P] ATP (~1 μCi) at 37°C for 30 min. Reactions were terminated by heating at 95°C for 3 min, and subsequently incubated with 4 U of alkaline phosphatase (New England Biolabs) at 37°C for 30 min to hydrolyze unreacted NTPs. 1 μL of each reaction was spotted on a PEI-Cellulose F thin-layer chromatography plate (EMD Biosciences) developed with 1.5 M KH_2_PO_4_ (pH 3.8) as a running buffer.

### Live imaging confocal microscopy

GFP-tagged cGAS-expressing iBMDMs were plated on uncoated 35 mm dishes (MatTek Corporation). Cells were treated with MitoTracker Deep Red (Invitrogen) in Opti-MEM (Gibco) and incubated at 37°C, 5% CO_2_ for 60 minutes. Cells were washed with PBS and imaged using a 63× oil immersion objective on the LSM 880 with Airyscan (Zeiss). Images were processed using ZEN software (Zeiss) and ImageJ (NIH). Dox-untreated cells or non-GFP-expressing cells were used for the negative controls of GFP signaling, and negative signals were subtracted from GFP signals of cGAS. For counting, at least 100 cGAS-expressing cells were examined under the microscope and the percentile of nuclear, cytosolic, or mitochondrial cGAS-expressing cells was determined.

### Quantification and statistical analysis

Statistical significance was determined by two-way analysis of variance (ANOVA) with Tukey’s multiple comparison test. *P* < 0.05 was seen as statistically significant. All statistical analyses were performed using GraphPad Prism data analysis software. All experiments were performed at least three times, and the graph data with error bars indicate the means with the standard error of the mean (SEM) of all repeated experiments.

## Supplementary Materials

**Materials and methods**

**Fig. S1 to S2**

Movie S1

## Acknowledgments

We thank all the members of the Kagan laboratory for helpful discussions.

## Funding

This work was supported by NIH grants AI133524, AI093589, AI116550 and P30DK34854 to J.C.K. D.C.H. is funded by 1R35GM142689-01 and a Recruitment of First-Time, Tenure-Track Faculty Award from the Cancer Prevention & Research Institute of Texas (RR 170047).

## Author contributions

K.M. designed the study, performed experiments, and wrote the manuscript. W.Z. and A.A.G. performed *in vitro* cGAMP analysis under the supervision of P.J.K. J.C.K. conceived the idea, supervised the research, and wrote the manuscript. D.C.H. provided critical reagents that supported these studies. All authors discussed the results and commented on the manuscript.

## Competing interests

J.C.K. consults for IFM Therapeutics and consults and holds equity in Corner Therapeutics, Larkspur Biosciences and Neumora Therapeutics. None of these relationships influenced the work performed in this study.

## Data and materials availability

All data needed to evaluate the conclusions in the paper are present in the paper or the Supplementary Materials. Further information and requests for resources and reagents should be directed to and will be fulfilled by the lead contact, J.C.K.

## SUPPLEMENTARY MATERIALS

### Supplementary Material and Methods

#### Generation of CRISPR-Cas9-mediated knockout (KO) cells

For cloning, the sense and antisense oligonucleotides containing guide RNA (gRNA) sequences and AfeI/SbfI restriction sites were purchased from Integrated DNA Technologies, and the equal molar ratios of sense and antisense oligonucleotides were annealed in water on PCR block (95°C for 1 minute, then drop 5°C every minute to 10°C). Oligonucleotide duplexes were then subcloned into AfeI/SbfI-digested pRRL-Cas9-Puro vector (kindly provided by Dr. D. Stetson (*8*)) using In-Fusion Snap Assembly Master Mix (TakaraBio). gRNAs targeting mouse genes used in this study are as follows: *Cgas* #1: GAGGCGCGGAAAGTCGTAAG, *Cgas* #2: GGCAGCCCAGAGCGCCGCGA, *Sting* #1: GGCCAGCCTGATGATCCTTT, *Sting* #2: GCTGGCCACCAGAAAGATGA.

To generate lentiviral particles for the stable expression of Cas9 and gRNAs, HEK293T cells were transfected with the packaging plasmids psPAX2 and pCMV-VSV-G along with the gRNAs-containing pRRL-Cas9-Puro vector using Lipofectamine 2000 (Invitrogen). Plasmids were transfected into 10 cm^2^ dishes of HEK293Ts at 50%–80% confluency using Lipofectamine 2000 by mixing DNA and Lipofectamine 2000 ratio at 1:2. Media was changed on transfected HEK293Ts 16-24 hours after transfection, and virus-containing supernatants were harvested 24 hours following the media change. Viral supernatants were passed through a 0.45 μm filter to remove any cellular debris. Filtered viral supernatants were mixed with polybrene (Millipore) and placed directly onto target cells, followed by the spinfection (centrifugation at 1,250 x g for 1 hour). Cell culture media were replaced with the appropriate complete media and cells were incubated for 24 hours. Spinfection with the viral supernatants was repeated on the following day, and cells were used for indicated assays.

#### Immunoblotting

Immunoblot analysis was performed as described in the main manuscript. For the detection of cGAS in Figure S1G, the primary antibody purchased from Cell Signaling Technology was used (1:1000 dilution).

**Fig. S1.**
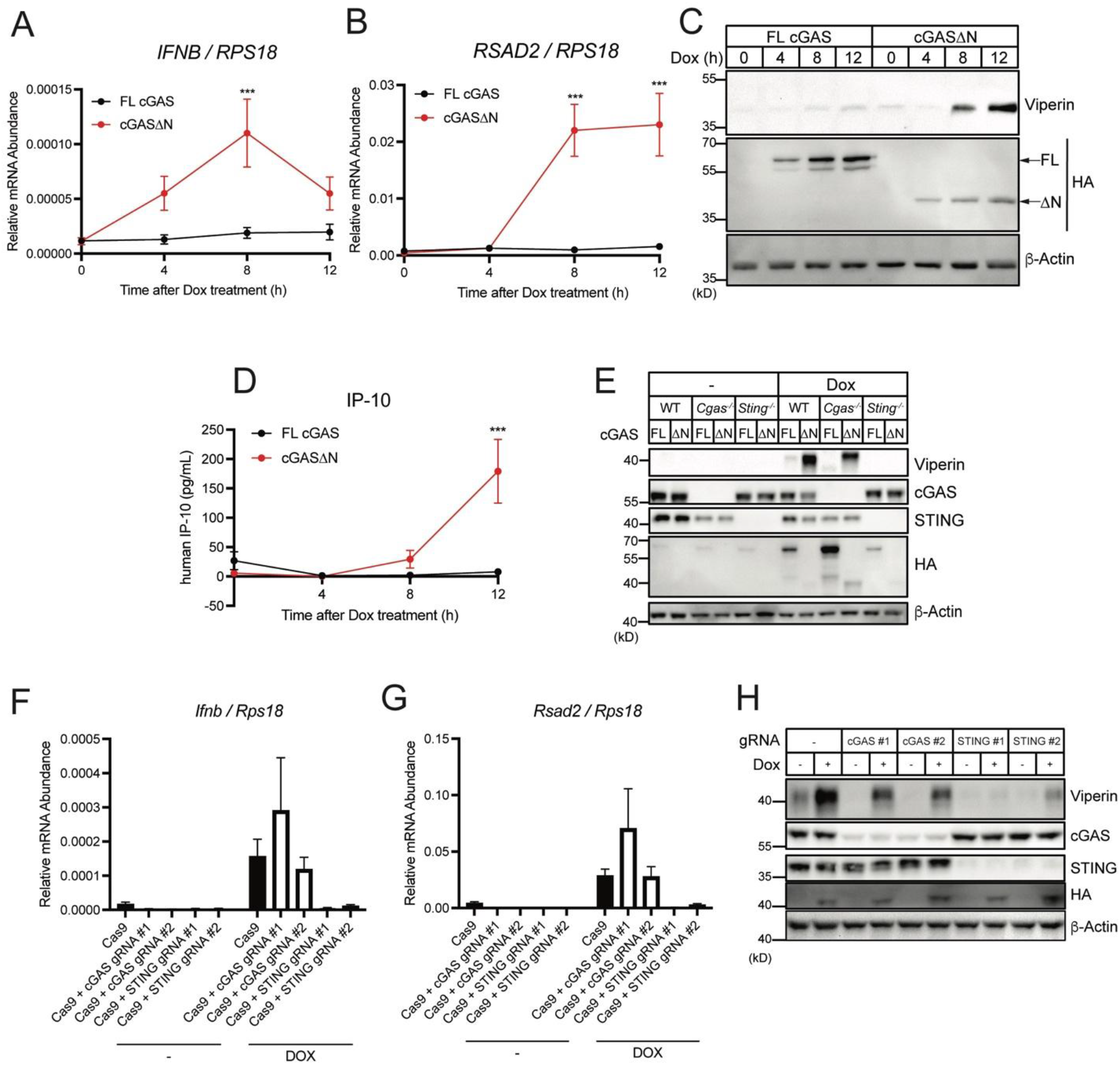
Human cGASΔN induces type I IFN responses. (**A** and **B**) Real-time qRT-PCR analysis of *IFNB* (A) and *RSAD2* (B) mRNAs in normal oral keratinocytes (NOKs). Dox was treated to cells for the induction of FL human cGAS or human cGASΔN, and mRNA expression levels were analyzed at the indicated time points. (**C**) Immunoblot analysis of NOK cell lysates after the same treatment as in (A) and (B). Arrows in the HA panel indicate FL human cGAS and human cGASΔN. (**D**) IP-10 ELISA analysis of NOK cell culture supernatant of the cells in (A) and (B). (**E**) Immunoblot analysis of lysates from WT, *Cgas*^-/-^, or *Sting*^-/-^ iBMDMs treated with Dox for 8 hours to induce FL human cGAS or human cGASΔN expression. (**F** and **G**) Real-time qRT-PCR analysis of *Ifnb* (F) and *Rsad2* (G) mRNAs in iBMDMs stably expressing Cas9 and indicated (guide) gRNAs treated with Dox for 8 hours for cGAS expression. (**H**) Immunoblot analysis of iBMDMs treated with Dox as in (F) and (G). Immunoblot data are the representative from three independent experiments. Graph data are means ± SEM of three (B), four (A), five (F and G), or six (D) independent experiments. Statistical significance was determined by two-way ANOVA and Tukey’s multicomparison test. Asterisks indicate the statistical significance between FL hcGAS and hcGASΔN at each time point (A, B, and D). \**P* < 0.05; \*\**P* < 0.01; \*\*\**P* < 0.001.

**Fig. S2.**
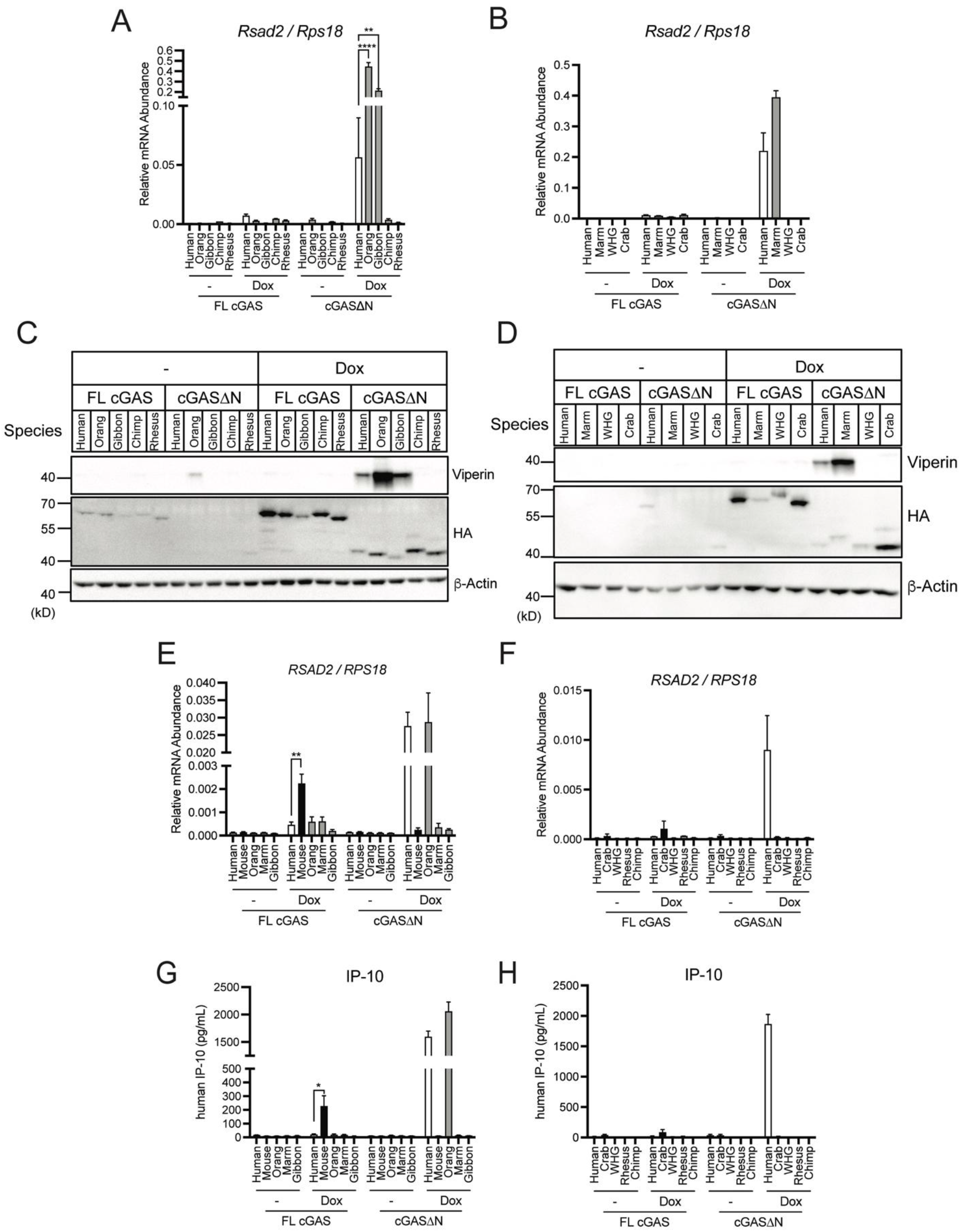
cGASΔN activities vary across mammalian species. (**A** and **B**) Real-time qRT-PCR analysis of *Rsad2* mRNA in iBMDMs treated with Dox for 8 hours to induce the expression of indicated cGAS. (**C** and **D**) Immunoblot analysis of iBMDM lysates treated as in (A) and (B). (**E** and **F**) Real-time qRT-PCR analysis of *RSAD2* mRNA in THP-1 cells treated with PMA and Dox for 48 hours to induce the expression of indicated cGAS. (**G** and **H**) IP-10 ELISA analysis of the THP-1 culture supernatant in (E) and (F). Immunoblot data are the representative from three independent experiments. Graph data are means ± SEM of three independent experiments. Statistical significance was determined by two-way ANOVA and Tukey’s multicomparison test. Asterisks indicate the statistical significance between connected two bars. \**P* < 0.05; \*\**P* < 0.01; \*\*\**P* < 0.001.

**Movie S1. Human cGASΔN localizes in the mitochondria.** A 3D reconstruction of z-stack images. Green: hcGASΔN-GFP, Red: Mitochondria. Images were processed using Zen software. The movie is representative from three independent experiments.

